# Metabolic demand regulates selective cell death in Parkinson’s Disease

**DOI:** 10.1101/2021.10.24.463836

**Authors:** Luke Zhao, Ishaan Nanal, Linxi Wang, Andrew Looka, Marianne Bezaire

**Affiliations:** RISE Neurobiology Program, Boston University, Boston MA 02215, USA; Lynbrook High School, San Jose CA 95129, USA; BASIS Independent Silicon Valley, San Jose CA 95126, USA; Canyon Crest Academy, San Diego CA 92130, USA; Boston University Medical Sciences Graduate Program, Boston University School of Medicine, Boston MA 02215, USA; Department of Psychological and Brain Sciences, Boston University, Boston MA 02215, USA

**Author notes:** These authors contributed equally to this work (Luke Zhao, Ishaan Nanal, Linxi Wang). Corresponding authors. or one of the first authors.

## Abstract

Parkinson’s Disease (PD) is a debilitating neurodegenerative condition that affects over 10 million people across the world, causing tremors and muscle weakness. Its mechanisms are unknown, but one key feature is selective cell death: neurons in the Substantia Nigra Pars Compacta (SNc) die, but their neighbors, the cells in the Ventral Tegmental Area (VTA), remain healthy. To study this phenomenon, we used an established single neuron model of the SNc, adapting its biophysical and bioenergetic properties to match that of the VTA. We discovered that reducing calcium influx correlates with higher ATP and lower ROS concentrations in the cell, suggesting in silico the importance of calcium influx in metabolic stress and selective vulnerability for Parkinson’s Disease. Future efforts may target calcium channel inhibition as a therapeutic strategy, although caution is needed with potential metabolic side effects.

## Introduction

At the moment, Parkinson’s disease is the second most common neurodegenerative disease in the world. While many therapies have been helping treat the disease, an absolute cure has yet to be identified, and until we know the exact cause of Parkinson’s, we will not be able to synthesize a cure. There have been a few theories postulating the underlying cause of Parkinson’s. One of the front-running theories argues for selectively high metabolic demands. This theory states that the reason behind SNc neuron vulnerability is their high metabolic demand. Since SNc cells are larger than other dopaminergic neurons in the brain, have more synaptic connections, and allow greater calcium influx when they release dopamine, they require more ATP to function. As a result, a small drop in ATP for an SNc cell causes a much bigger ripple effect than it would for other neurons. Neighboring the SNc, the VTA is an area of the brain that also contains dopaminergic neurons but is unaffected in Parkinson’s. By modeling both VTA and SNc cells and examining the differences in their biophysical properties, we can offer proof for the theory that SNc cells are selectively vulnerable to Parkinson’s due to their high metabolic demands.

## Methods

In order to test our hypothesis we would need two model codes. One that would model a VTA neuron and one that would model a SNc neuron. Our models would need to simulate both the biophysical properties of a cell as well as its metabolic pathways. We were able to utilize the Substantia Nigra pars Compacta model created by Vignayanandam Ravindernath Muddapu & V. Srinivasa Chakravarthy from the Indian Institute of Technology Madras^[1]^. This model was created on MATLAB version R2020a, and provided us with the simulation of a SNc neuron. In order to simulate a Ventral Tegmental Area neuron we took the model of the SNc neuron and changed the electrophysical properties in the code to better simulate a VTA neuron. The first parameter that was changed was the resting membrane potential. The resting membrane potential of a SNc neuron is -49.42 mV. The resting membrane potential of a VTA neuron is -51.69 mV, which is indicative of the difference in cellular properties. By changing this parameter, the electrophysiological properties were changed from that of the Substantia Nigra pars Compacta to that of the Ventral Tegmental Area. In addition, while SNc neurons and VTA neurons are both dopaminergic they differ in their subcellular processes and ion concentrations, specifically Ca^2+[2]^. The maximal conductance of the calcium channel of the SNc neuron was 2101.2 pA/mM. This was changed by a scaling factor of 1.00 to 0.10 to better replicate the lower Ca^2+^ ion influx in VTA neurons.

### Development of a VTA neuron model

SNc and VTA neurons are highly similar in function and biological pathways. To model the SNc neurons, we used a previously created model of SNc bioenergetics and Parkinson’s pathways^1^. Model results are shown (Figure 1). To adapt our model of the SNc neuron for a VTA neuron, we altered two key parameters: resting membrane potential and maximal calcium ion channel conductance. Resting membrane potential is a baseline cell property that varies from neuron to neuron, providing an anchoring feature for the VTA neuron. From the database NeuroElectro, which provides neuron electrophysiological properties, we used the averaged dopaminergic VTA resting membrane potential of -51.69 mV. This differs from the original SNc resting membrane potential of -49.42 mV (Figure 2). Next, we reduced maximal conductance of the L-type calcium channel (G_Ca_) by a scaling factor between 0 and 1, as done previously^2^. Maximal conductance is a measure of how easily an electrical current can pass through the channel. By reducing maximal conductance in our model, we directly reduced the maximum calcium influx through these ion channels, reflecting the lower calcium influx seen in VTA neurons in vivo (Figure 3).

**Figure 1.**
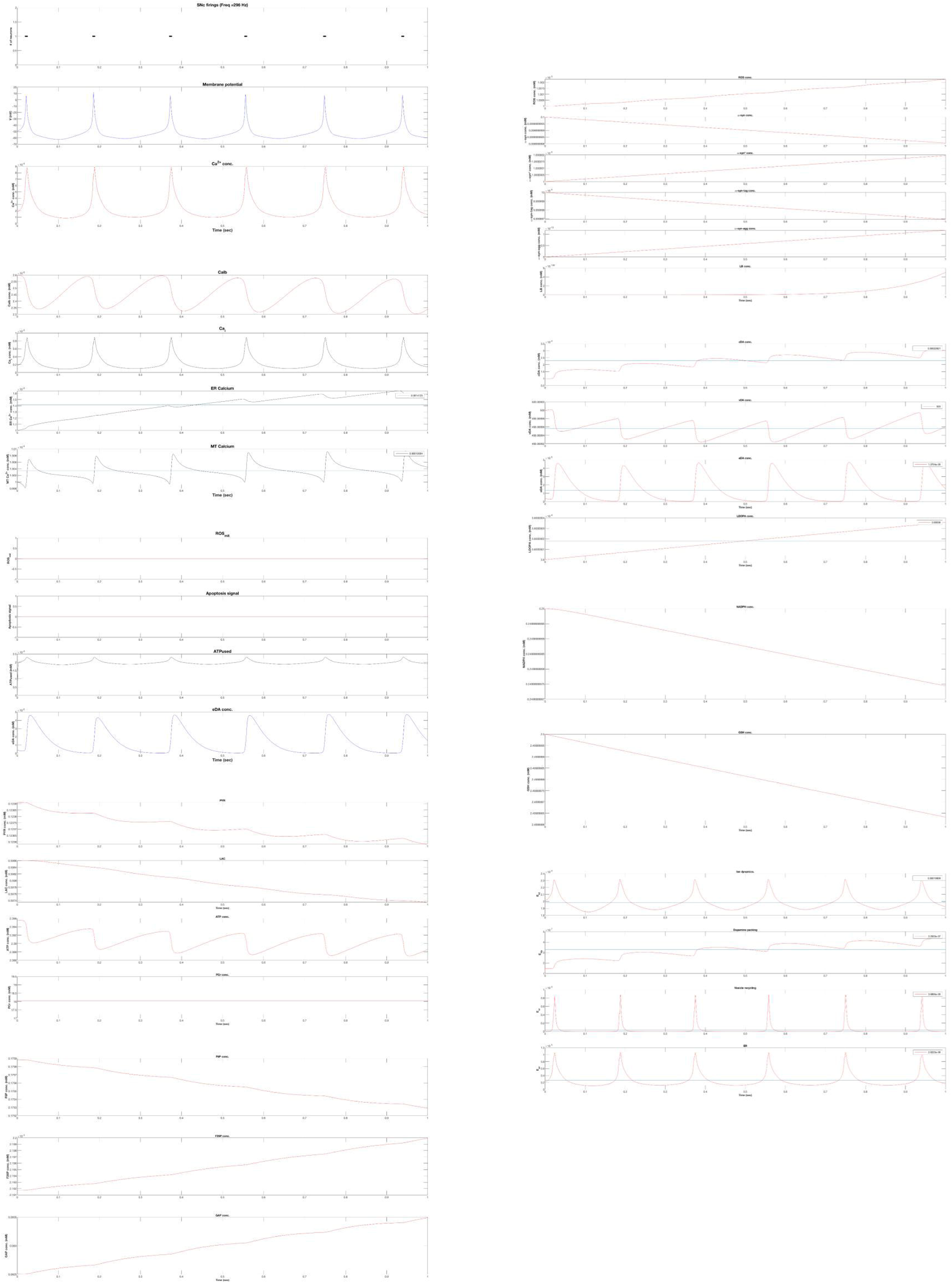
Modeled properties of the SNc neuron. Properties include membrane voltage, calcium concentration, reactive oxygen species concentration, apoptosis signaling, ATP usage, extracellular dopamine concentration, metabolite concentrations, and others.

**Figure 2.**
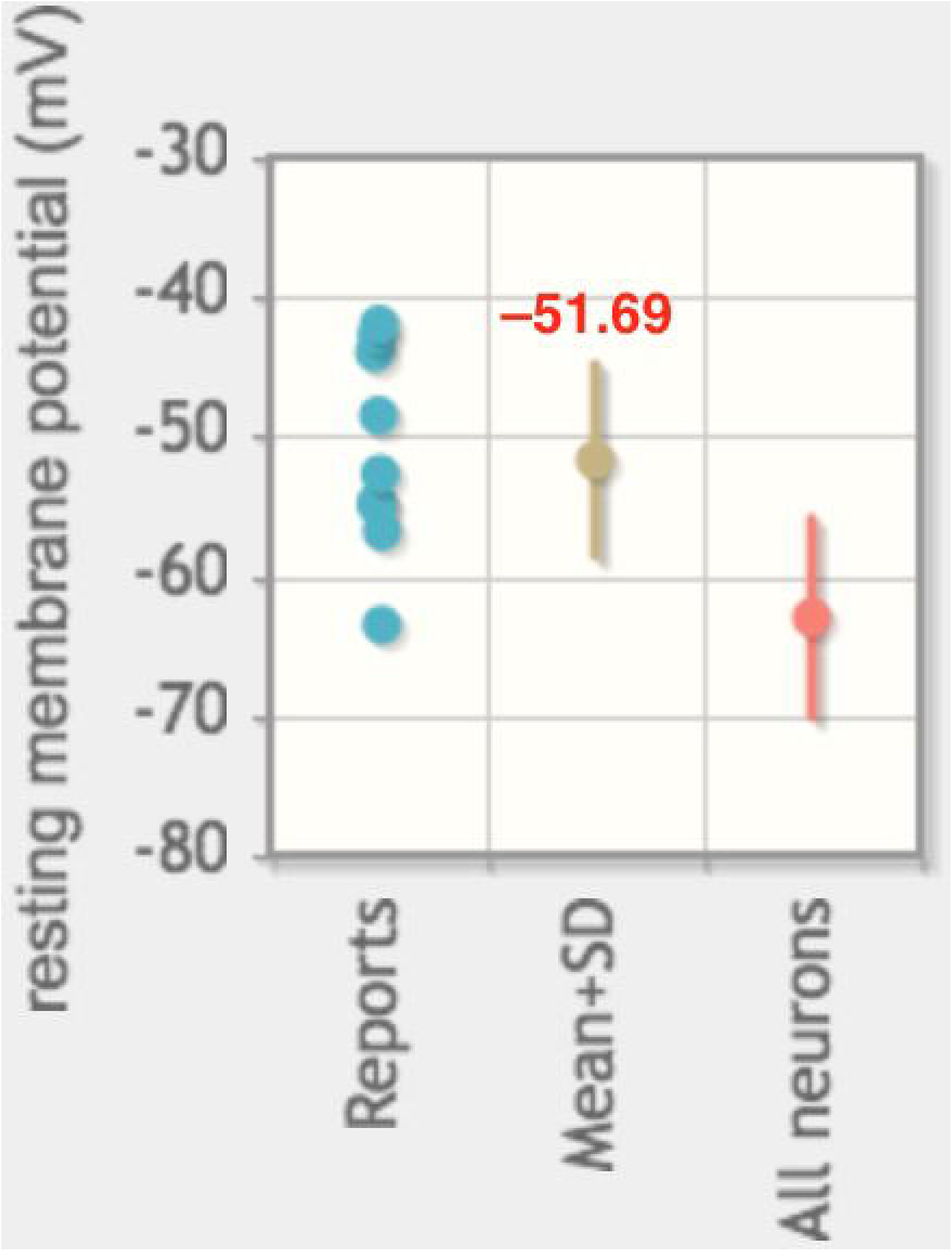
Resting membrane potential. Average resting membrane potential of VTA neurons from NeuroElectro

**Figure 3.**
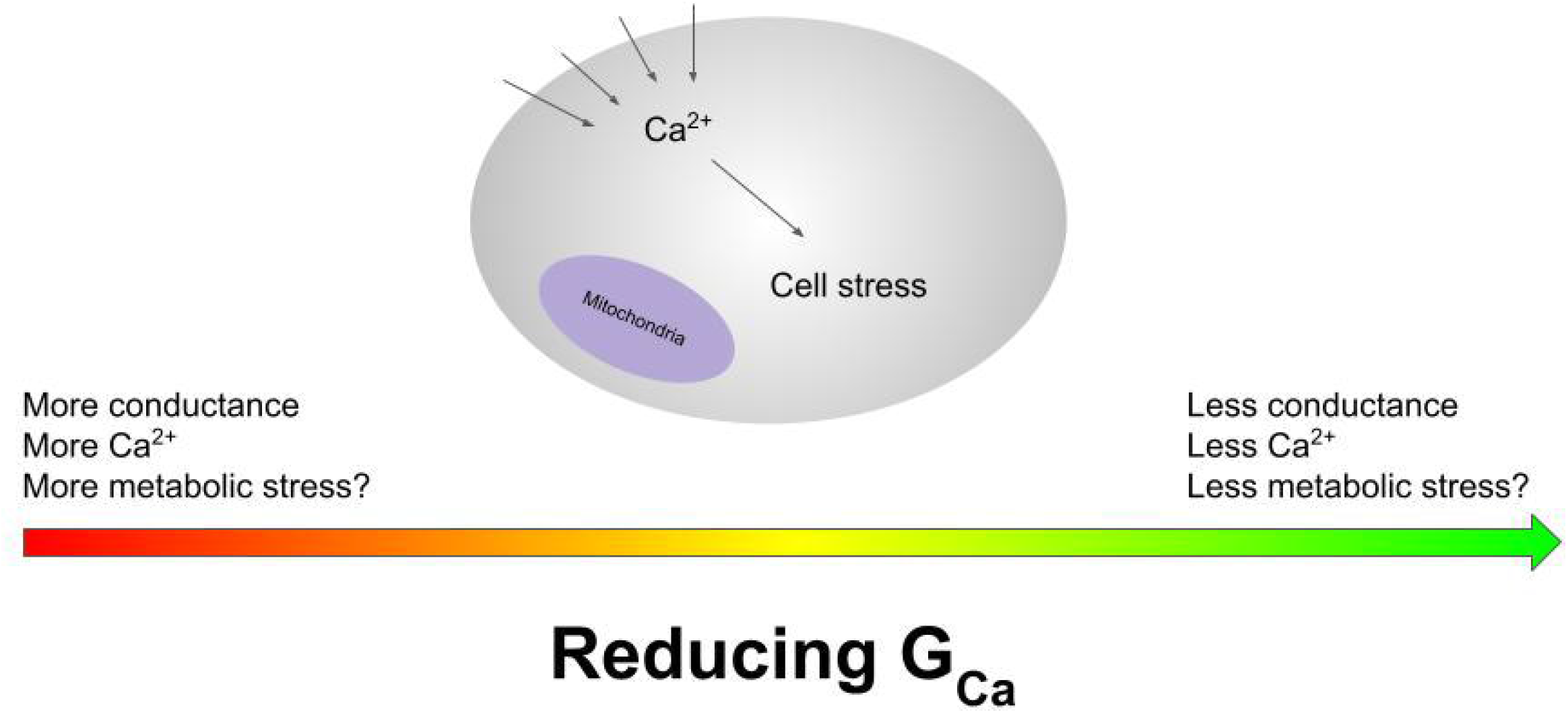
Proposed model. Reducing calcium conductance leads to reduced baseline metabolic stress

### Reduction of calcium channel maximal conductance

We reduced calcium channel conductance by five factors: 1.0x, 0.75x, 0.50x, 0.25x, and 0.10x. VTA neurons are recapitulated in each, but the degree of calcium influx is gradually reduced. This additionally models the effects of a calcium channel inhibitor, beyond the baseline lower calcium influx in VTA neurons. We observed changes in firing patterns, calcium dynamics, dopamine production, and baseline metabolic stress, as encoded by the model (Figures 4-11).

**Figure 4.**
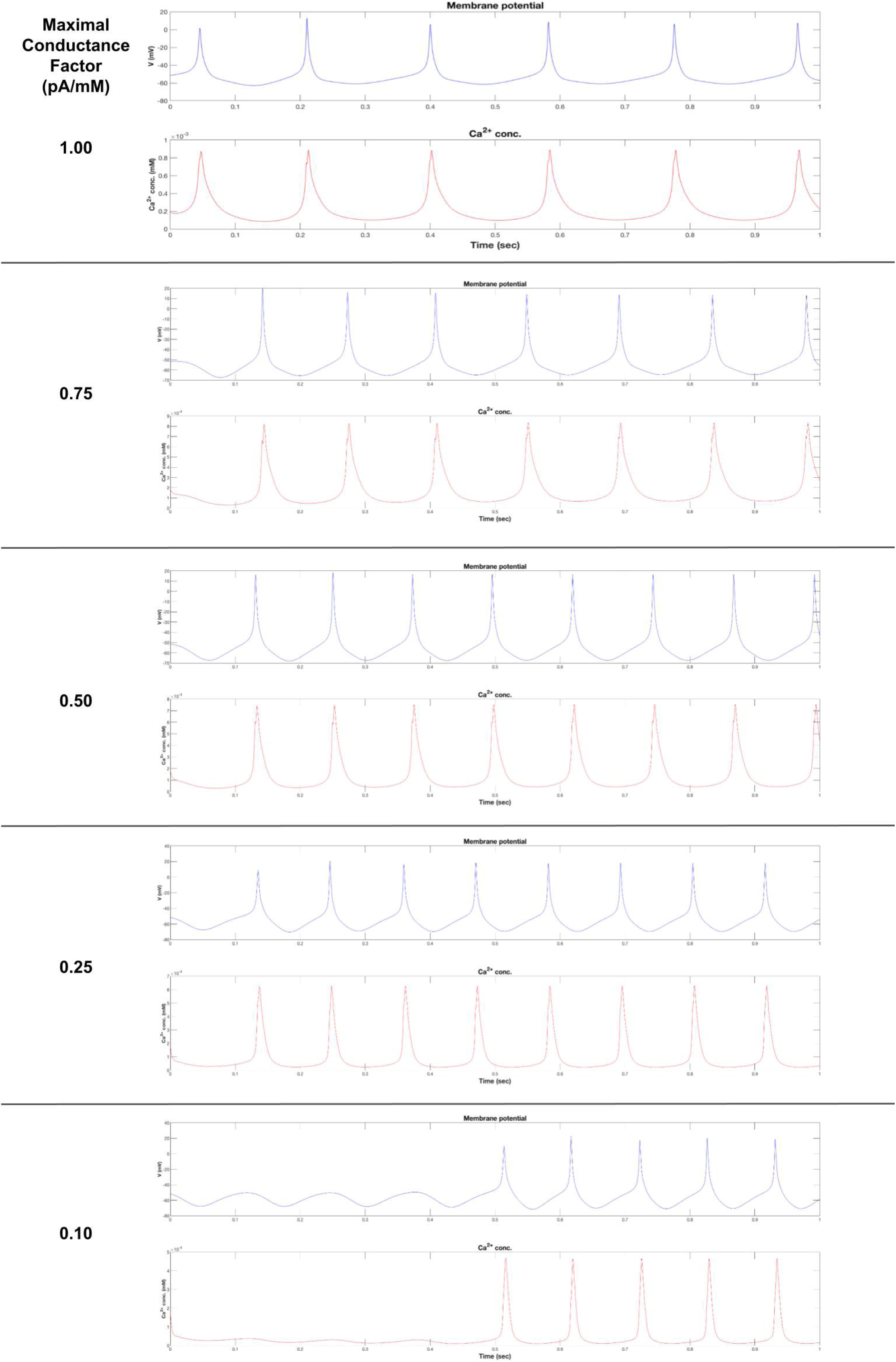
Reducing calcium conductance alters membrane potential and calcium concentration. (a) As calcium conductance is gradually decreased, spikes in membrane potential reach a greater maximum, near +10 mV. Continued decreases in conductance lead to tapered beginnings, representing a time in which the model adapts to decreased calcium influx. The relatively consistent firing also suggests the baseline functions of neurons are preserved, and the model recapitulates VTA neuron mechanics. (b) Calcium concentration is reduced significantly, and maximum spikes are lowered from 9×10^-4 to 5 × 10^-4 mM. This is expected since reducing conductance reduces calcium influx, so there should be less calcium in the neuron models. Calcium concentration also experiences a tapered beginning.

**Figure 5.**
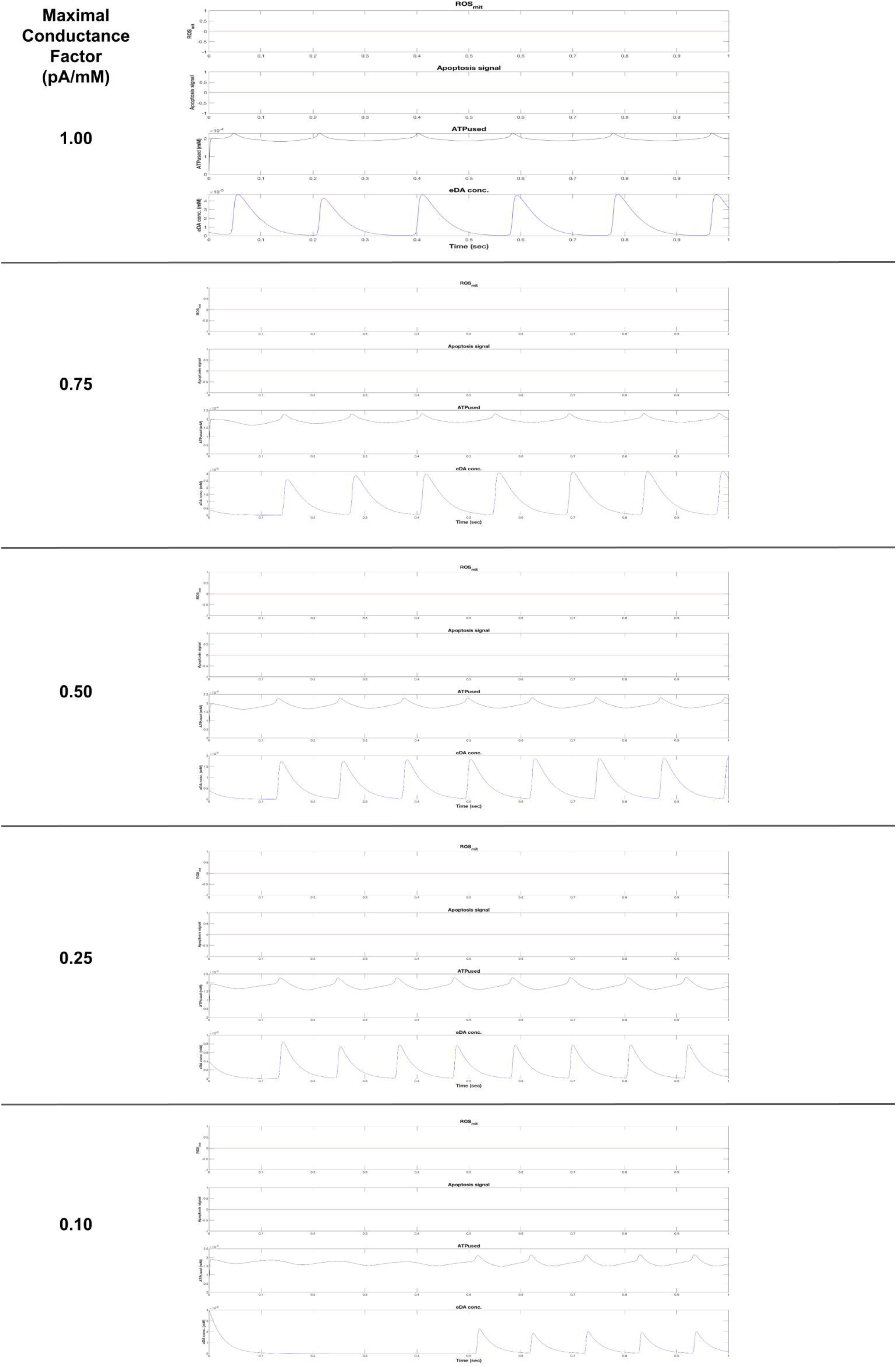
Reducing calcium conductance alters extracellular dopamine concentration. (a) Reactive oxygen species in the mitochondria are kept at a baseline level (no stress induced) (b) Apoptosis signaling is also kept at a baseline level, as expected (no stress induced) (c) ATP used is kept consistent. Regular spikes of ATP usage are observed, correlating with the regular firing patterns of neurons. (d) Extracellular dopamine concentration is significantly decreased. This is expected, since neurons depend on calcium signaling to release dopamine vesicles into synapses. The decrease warns that reducing calcium influx, by inhibitors or by adapting to a VTA neuron, leads to less dopaminergic function.

**Figure 6.**
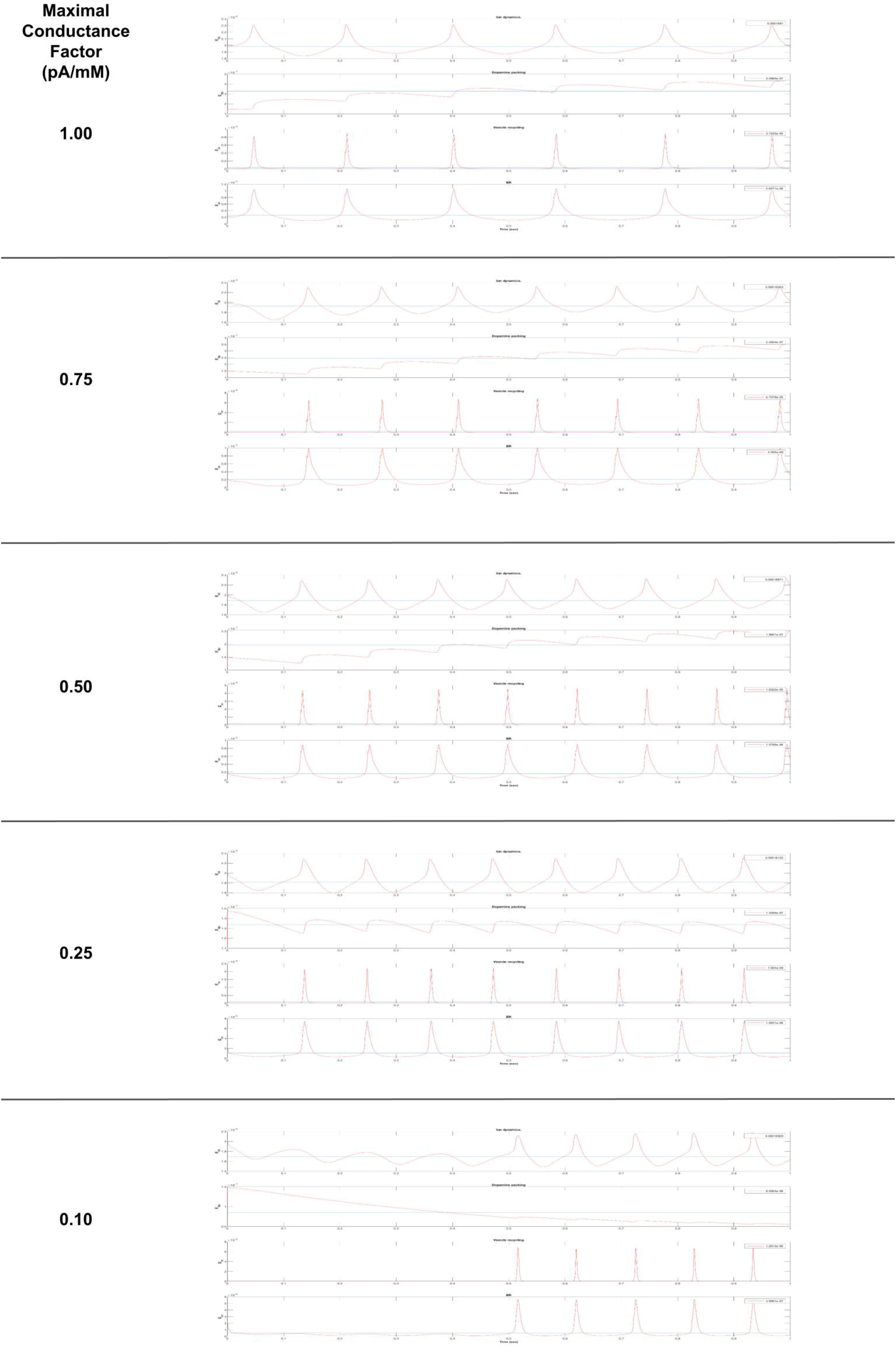
Reducing calcium conductance alters calbindin concentration, internal calcium concentration, ER calcium concentration, and mitochondrial calcium concentration. (a) Calbindin concentrations rise as calcium conductance is decreased. Possibly, the model adapts to lower calcium influx by increasing calbindin, attempting to buffer and regulate calcium concentrations. (b) Internal calcium is lowered greatly. This validates our proposed method of decreasing calcium influx computationally, and we observe a direct decrease in internal calcium. Calcium concentrations also spike regularly, as expected in a neuron. (c) Endoplasmic reticulum calcium concentrations are decreased, as expected. (d) Mitochondrial calcium concentrations are affected interestingly. Whereas a regular spiking pattern is observed with full calcium conductance, a decrease with tapered spikes is observed with 50% conductance and beyond.

**Figure 7.**
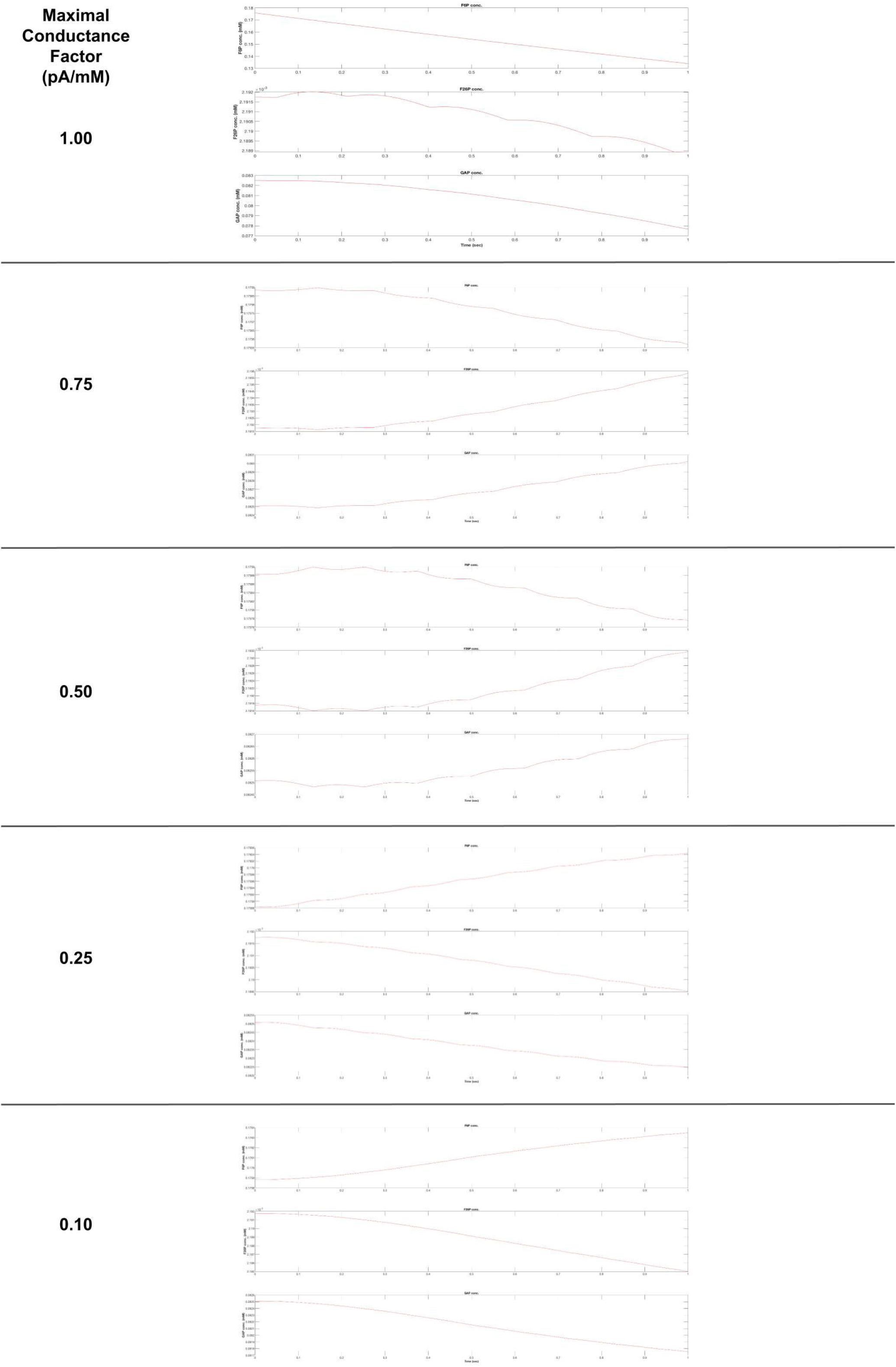
Reducing calcium conductance alters metabolite concentrations. (a) Fructose 6-phosphate (F6P) concentrations lower over time but then increase at and beyond 25% channel conductance. Possibly, not enough calcium is supplied beyond 25% conductance to allow normal mitochondrial functioning, resulting in an excess of glycolytic products and less need for F6P production. (b) Fructose 2,6-bisphosphate (F26P) concentrations lower over time but increase only for 75% and 50% channel conductance. Since F26P is a downstream glycolytic product of F6P, this is an artifact of F6P concentrations being affected earlier. (c) Glyceraldehyde-3-phosphate (GAP) concentrations lower over time but increase only for 75% and 50% channel conductance. Since GAP is a downstream glycolytic product of F6P, this is an artifact of F6P concentrations being affected earlier.

**Figure 8.**
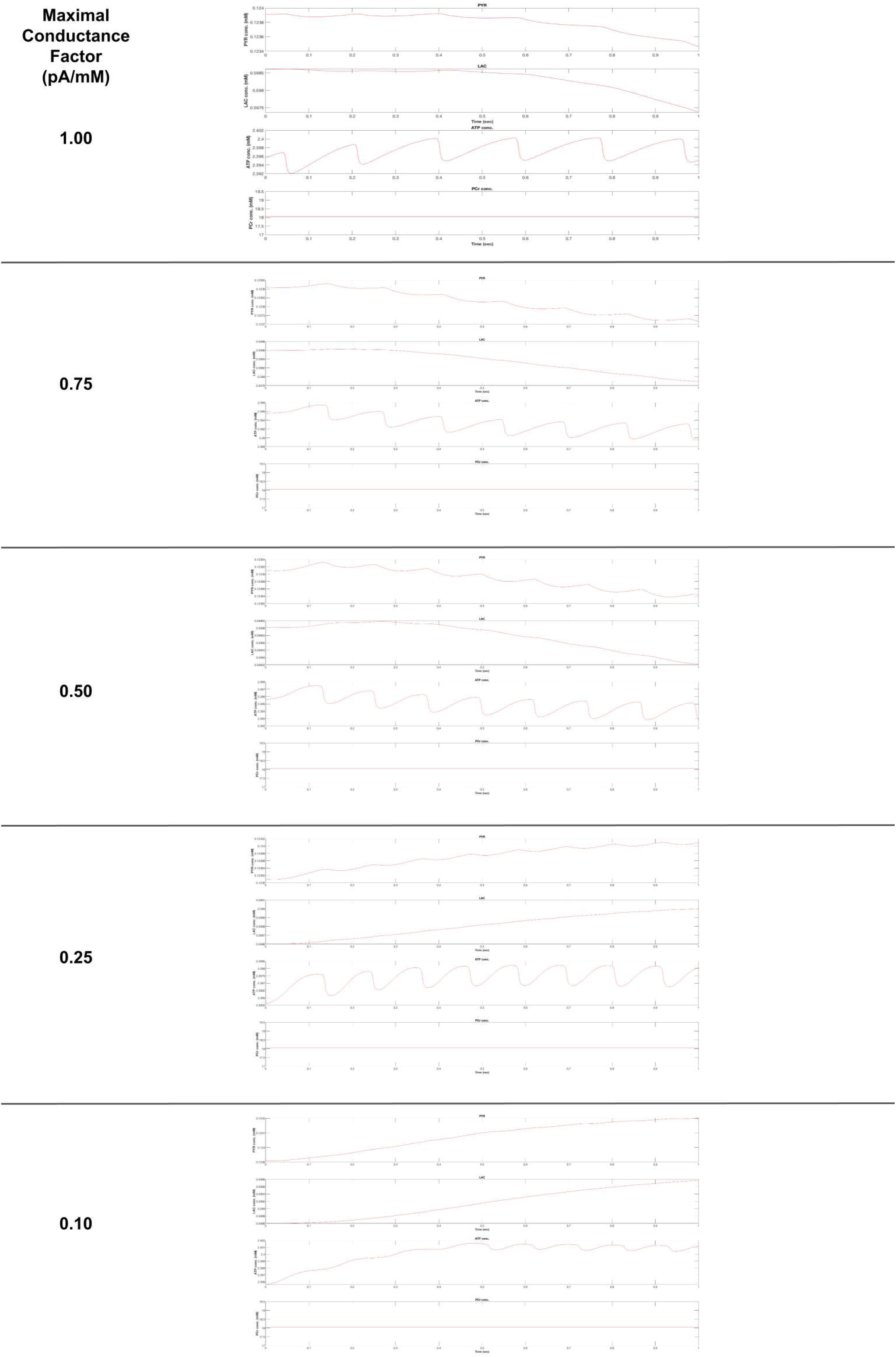
Reducing calcium conductance alters more metabolite concentrations, including ATP. (a) Pyruvate (PYR) concentrations lower over time but then increase at and beyond 25% channel conductance. Possibly, not enough calcium is supplied beyond 25% conductance to allow normal mitochondrial functioning, resulting in an excess of glycolytic products and less need for PYR production. Glycolysis outpaces mitochondrial activity. (b) Lactate (LAC) concentrations decrease over time from a baseline value, but concentrations increase at and beyond 25% channel conductance. This suggests mitochondrial functioning is impaired with 25% channel conductance, leading to the use of anaerobic respiration and the production of lactate. (c) Baseline 100% channel conductance ATP concentrations are generally consistent, fluctuating around 2.396 mM. At 75% channel conductance, there is a slight decrease, fluctuating around 2.392 mM. At 50% channel conductance, ATP fluctuates at 2.394 mM. At 25%, 2.3975 mM. At 10%, 2.41 mM. The initial decrease in ATP could suggest a model adaptation to lowered calcium influx. But as calcium influx is decreased further, ATP concentrations rise significantly, suggesting lower baseline metabolic stress. Less calcium enters the cell, so the cell uses less ATP in restoring ion equilibrium. Therefore, reducing calcium ion influx is a potential way to alleviate cell metabolic demand. VTA neurons, with less calcium influx, experience less baseline metabolic stress and are more resistant to Parkinson’s Disease. SNc neurons on the other hand, with more calcium influx, experience more metabolic stress and degenerate in Parkinson’s. Reducing calcium ion influx can be a potential therapeutic method to alleviate Parkinson’s in SNc neurons. (d) Phosphocreatine (PCr) concentrations remain constant, indicating these pathways of metabolism are not used in our model.

**Figure 9.**
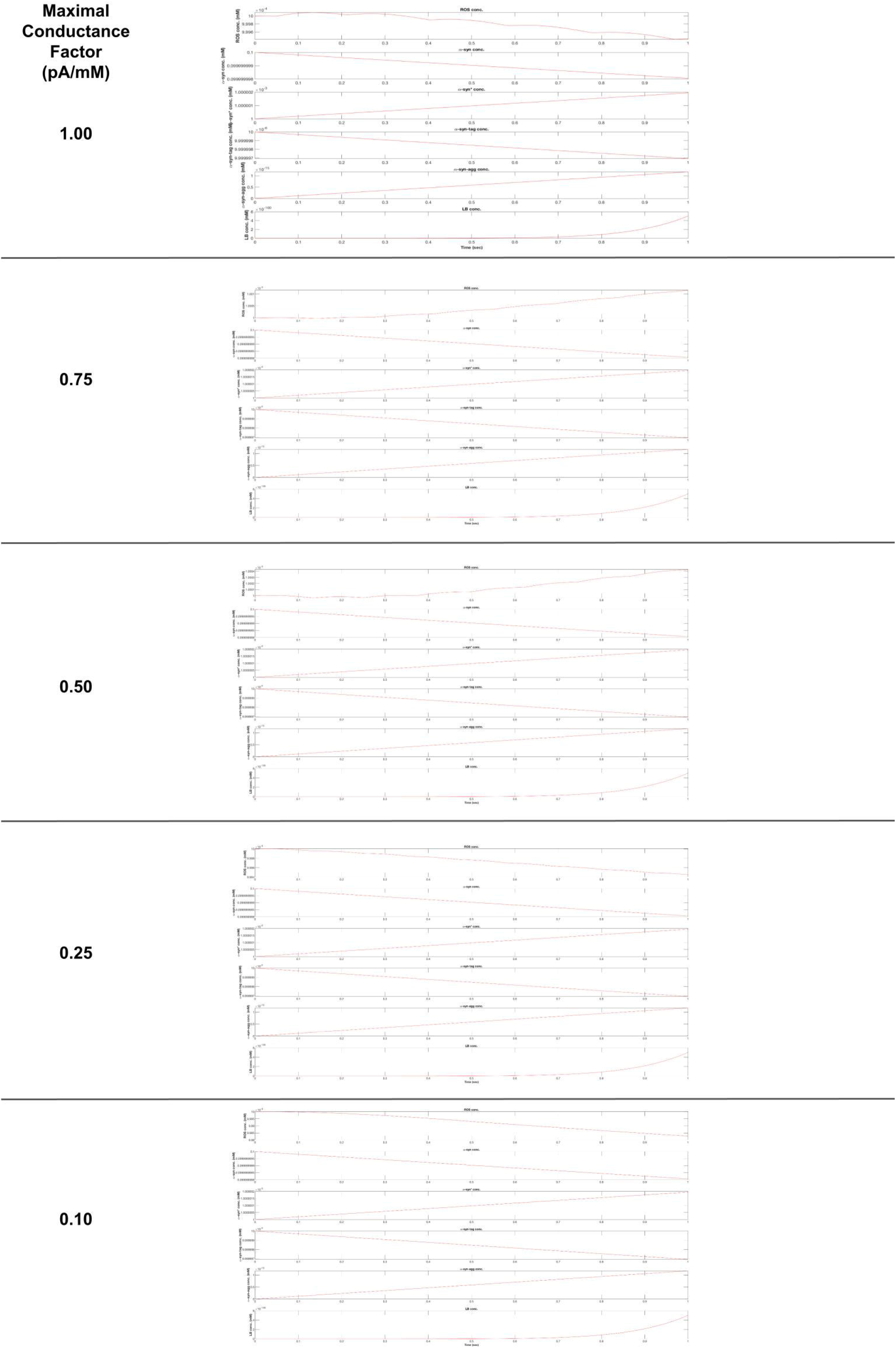
Reducing calcium concentration alters ROS concentration. (a) Reactive Oxygen Species (ROS) starts from a baseline concentration of 1×10^-4 mM. With 100% channel conductance, ROS decreases to 0.9994 × 10^-4 mM over the model run time (1000 ms). With 75%, ROS increases to 1.001×10^-4 mM. With 50%, ROS increases to 1.0004×10^-4 mM. With 25%, ROS decreases to 0.9994 × 10^-4 mM. With 10%, ROS decreases to 0.9985×10^-4 mM. Conclusively, ROS concentrations were decreased past the baseline reduction with 25% channel conductance. (b) Alpha-synuclein concentration is unaffected. (c) Misfolded alpha-synuclein concentration is unaffected. (d) Tagged alpha-synuclein concentration is unaffected. (e) Aggregated alpha-synuclein concentration is unaffected. (f) Lewy Body concentration is unaffected.

**Figure 10.**
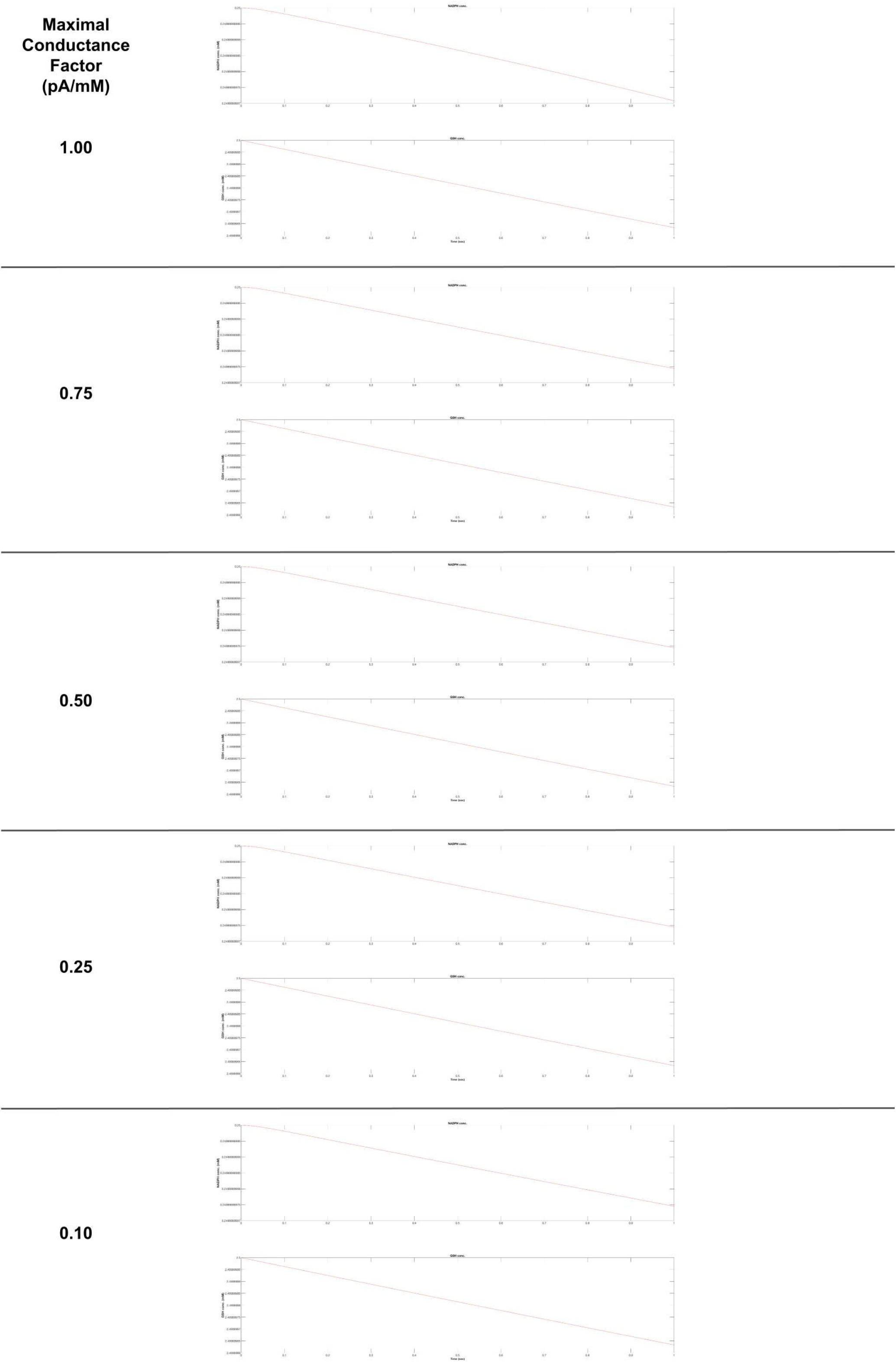
Reducing calcium conductance does not affect NADPH and GSH concentration. (a) NADPH concentration is unaffected. Parallel to glycolysis, F6P is used to produce NADPH by the pentose phosphate pathway. NADPH is used by glutathione reductase to synthesize glutathione (GSH). (b) GSH concentration is unaffected. GSH is a reducing agent and an antioxidant, scavenging reactive oxygen species from the cell.

**Figure 11.**
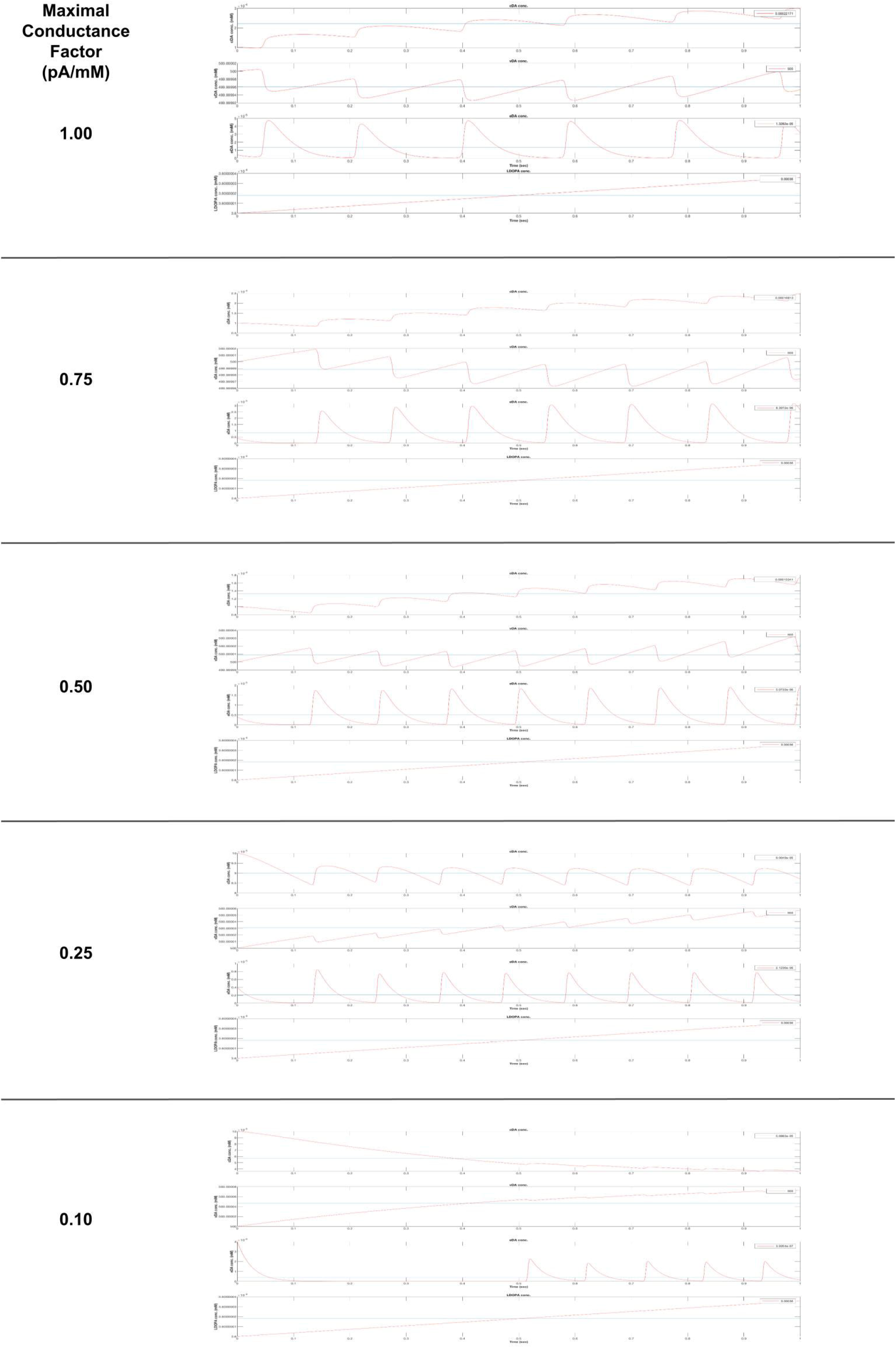
Reducing calcium conductance alters cytoplasmic dopamine, vesicular dopamine, extracellular dopamine, and L-DOPA production. (a) Average cytoplasmic dopamine (cDA) decreases as ion conductance is reduced. The average concentration (blue line) drops from 2.2×10^-4 mM to 0.59×10^-4 mM. (b) Vesicular dopamine (vDA) concentration follows a fluctuating pattern, but it loses this fluctuating pattern and average values increase marginally as ion conductance is reduced. (c) Extracellular dopamine (eDA) follows the same pattern as described in figure 4. (d) L-DOPA concentration is unaffected. L-DOPA is a dopamine precursor molecule.

## Discussion

Our results demonstrate that reducing calcium conductance is beneficial to cell survival and metabolic stress. We highlight two key factors: ATP concentration and ROS concentration. We observed that as the scaling factor was reduced from 1.00 to 0.1, the ATP concentration in the VTA neuron increased. This would indicate that as less calcium was allowed into the cell, the ATP concentration within the cell increased. Because ATP is increased, the cell has higher metabolic flexibility at a baseline level. As the scaling factor decreased in the VTA neuron, the ROS concentration also decreased. ROS concentration is an indicator of cell metabolic stress, so the reduced ROS concentration is a positive, anti-stress effect of reducing calcium conductance.

Some results were unexpected. For example, alpha-synuclein and Lewy Body concentrations were unaffected by reducing calcium conductance. We expected these to decrease, as ROS concentration did. Possibly this is how the model code is originally designed, where avoiding changes in glucose and oxygen concentrations initially will not trigger downstream Parkinson’s-specific pathology. Additionally, NADPH and GSH concentrations were unaffected. We expected these to increase since glutathione scavenges reactive oxygen species, but the drop in ROS concentration could have been caused by lower mitochondrial functioning or other changes as well. Likewise, our proposed method of reducing calcium channel conductance models the effects of a calcium channel inhibitor in silico. However, reduction of conductance at and beyond 25% of its original functionality had side effects to the neuron model. Dopamine production and release into the extracellular space was markedly decreased. This suggests future attempts to inhibit calcium channels in Parkinson’s may decrease the dopaminergic function of SNc cells –a consideration that needs to be made when considering drug design. Additionally, mitochondrial function was impaired. Production of glycolytic metabolites like F6P and GAP were significantly increased while concentrations of lactate were also increased, suggesting the mitochondria depend on sufficient calcium influx to perform their normal functions. When calcium influx is inhibited too severely, as in 25% channel functionality, cells resort to anaerobic respiration, produce lactic acid, and accumulate glycolytic byproducts in their cytoplasms. There are other limitations to our model: the extracellular environment for the VTA neuron isn’t exact, and remodeling the environment entirely will help achieve more precise results. In addition, getting in vivo confirmation for the theory would be best, since that would bypass the limitation of modeling in general. After confirming our findings through a more precise model, or through in vivo confirmation, a drug could be developed that inhibits calcium channels in the SNc neurons, with correct inhibitory efficacy. This would serve as a possibly more effective therapy for Parkinson’s Disease.

## Conclusion

We adapted a single neuron model of the SNc to represent the VTA neuron, based on changes in resting membrane potential and calcium channel conductance. Using our models of the SNc and VTA, we investigated differences in metabolic demand that lead to Parkinson’s Disease. With lower calcium channel conductance, we found ATP concentration increased and ROS decreased, indicating calcium influx plays a key role in metabolic susceptibility to Parkinson’s. Further inhibition of calcium channels may lead to greater results, but consideration is needed with the side effects of channel inhibition. Finally, future work is needed to confirm these results and elaborate on the role of calcium influx in Parkinsonian degeneration.

## Notes

### Competing Interest Statement

The authors have declared no competing interest.

### Summary of Updates

Manuscript formatting revised, author contributions updated (all contributed equally)

